# The potential for a released autosomal X-shredder becoming a driving-Y chromosome and invasively suppressing wild populations of malaria mosquitoes

**DOI:** 10.1101/860551

**Authors:** Yehonatan Alcalay, Silke Fuchs, Roberto Galizi, Federica Bernardini, Roya Elaine Haghighat-Khah, Douglas B. Rusch, Jeffrey R. Adrion, Matthew W. Hahn, Pablo Tortosa, Philippos Aris Papathanos

**Affiliations:** Department of Entomology, Robert H. Smith Faculty of Agriculture, Food and Environment, Hebrew University of Jerusalem, Rehovot 7610001, Israel; Department of Life Sciences, Imperial College London, London, United Kingdom; Center for Genomics and Bioinformatics, Indiana University, Bloomington, Indiana 47405, USA; Institute of Ecology and Evolution, University of Oregon, Eugene, OR 97403; Department of Biology, Indiana University, Bloomington, IN 47405; Department of Computer Science, Indiana University, Bloomington, IN 47505; Unité Mixte de Recherche Processus Infectieux en Milieu Insulaire Tropical (UMR PIMIT), Université de La Réunion, INSERM 1187, CNRS 9192, IRD 249, Plateforme de Recherche CYROI, Sainte-Clotilde, Reunion, France

## Abstract

Synthetic sex-ratio distorters based on X-chromosome shredding are predicted to be more efficient than sterile males for population suppression of malaria mosquitoes using genetic control. X-chromosome shredding operates through the targeted elimination of X-chromosome-bearing gametes during male spermatogenesis, resulting in males that have a high fraction of male offspring. Strains harboring autosomal constructs containing a modified endonuclease I-*Ppo*I have now been developed in the malaria mosquito *Anopheles gambiae*, resulting in strong sex-ratio distortion towards males. Data are being gathered for these strains for submission of regulatory dossiers for contained use and subsequent field release in West Africa. Since autosomal X-shredders are transmitted in a Mendelian fashion and can be selected against their frequency in the population is expected to decline once releases are halted. However, any unintended transfer of the X-shredder to the Y-chromosome could theoretically change these dynamics: This could lead to 100% transmission of the newly Y-linked X-shredder to the predominant male-biased offspring and its insulation from negative selection in females, resulting in its potential spread in the population and ultimately to suppression. Here, we analyze plausible mechanisms whereby an autosomal X-shredder could become linked to the Y-chromosome after release and provide data regarding its potential for activity should it become linked to the Y-chromosome. Our results strongly suggest that Y-chromosome linkage through remobilization of the transposon used for the initial genetic transformation is unlikely, and that, in the unexpected event that the X-shredder becomes linked to the Y-chromosome, expression and activity of the X-shredder would likely be inhibited by meiotic sex chromosome inactivation. We conclude that a functioning X-shredding-based Y-drive resulting from a naturally induced transposition or translocation of the transgene onto the Y-chromosome is unlikely.

## Introduction

Mosquito species of the *Anopheles gambiae* complex are the main vectors of human malaria and pose an enormous burden on global health and economies. Every year, 300–500 million people are infected with malaria and approximately half a million die as a result of these parasitic infections, 70% of these being children less than 5 years old [1]. The progressive spread of insecticide resistant insects [2,3], combined with the current lack of an efficient malaria vaccine, has prompted the development of new methods. One of the most promising is genetic control, based on the release of laboratory-modified insects into the environment; these individuals mate with wild individuals and transmit control traits that can suppress or modify the targeted population[4,5]. The most commonly used approach to genetically control insects has been the mass release of sterile males — the so-called Sterile Insect Technique (SIT) [6,7]. Through mating with released sterile males, wild females are effectively sterilized, if they only mate once during their lifetime, because offspring of sterile males are incapable of development. If sufficient numbers of sterile males are released over a long enough period, the wild population can be effectively suppressed or even eradicated. However, the economic costs associated with mass production of sterile males, the difficulty of covering very large areas, combined with the need to maintain sterile releases for continued suppression, have restricted the implementation of this method to date.

One way to improve the efficiency of such approaches is through the release of fertile males that are daughterless. Since male mosquitoes do not contribute to disease transmission, releasing males that have viable and fertile sons can help to temporarily maintain the control trait in the population. Two strategies based on such fertile males have been developed in mosquitoes: fs-RIDL (for female-specific Release of Insects carrying Dominant Lethals) and sex ratio distorters based on X-chromosome shredding. fs-RIDL is based on a construct that is lethal to females that inherit it, so that daughters of released transgenic males born in the field and inheriting the transgene die before maturing, but sons survive and pass the transgene to their offspring [8,9]. Sex-ratio distortion based on X-chromosome shredding instead relies on the expression of a sequence-specific endonuclease during male spermatogenesis that recognizes and cleaves sequences that are both specific and abundant on the X-chromosome [10,11]. As a result, X-chromosome-bearing gametes are excluded from the fertilizing-sperm population, biasing offspring sex-ratios towards males [12–14]. Mathematical models suggest that both approaches can be more efficient than SIT in terms of the number of modified males that need to be released to achieve a similar level of population suppression [15,16]. Despite being more efficient, both fs-RIDL and autosomal X-shredders (where the transgene is located on an autosome) are self-limiting—i.e. the constructs underlying the phenotype will not spread in the population—because they are inherited in a Mendelian fashion and do not provide any fitness advantage over the wild type, unlike self-sustaining approaches such as those incorporating gene drive constructs [17,18]. Of the two male daughterless systems, autosomal X-shredders are predicted to be more efficient because daughters are effectively replaced with male-forming gametes pre-zygotically. Therefore, on average, twice the number of transgenic male descendants are produced per released male, increasing the effective number of modified males compared to fs-RIDL males. X-shredding also has the potential to be used for self-sustaining genetic control applications, in the form of Y-chromosome drive as originally proposed by Hamilton [5]. This could be done by linking a functional X-shredder to the Y-chromosome, in which case both the Y-chromosome and the X-shredder gain a transmission advantage through preferential inheritance of male-forming gametes [19].

An autosomal X-shredding sex-distorter was first developed in *An. gambiae* by Galizi et al. [11]. They used variants of the I-*Ppo*I endonuclease that specifically cut a specific DNA target sequence within the 28S ribosomal DNA locus, which in *An. gambiae* is located exclusively on the X chromosome in approximately 200-400 copies [20]. The I-*Ppo*I protein was modified to restrict its activity to spermatogenesis, since expression of the native I-*Ppo*I resulted in male sterility through postcopulatory, embryonic carryover and consequent dominant embryonic arrest due to the shredding of the maternally inherited X-chromosome [21]. These I-*Ppo*I variants were fused to eGFP and driven by the *An. gambiae beta-2 tubulin* regulatory regions, which become active in primary spermatocytes entering male meiosis [22]. The resulting transformation constructs also included the *DsRed* transformation marker driven by the neuron-specific *3xP3* promoter, and the entire cassette was flanked by piggyBac-specific left and right arms containing the inverted terminal repeat sequences (ITRs) (Figure 1A). These ITRs are characteristic for transposable elements and are required for recognition by the co-injected piggyBac (PB) transposase that catalyzes integration of the construct in the *An. gambiae* genome at TTAA sites. A number of transgenic strains carrying independent autosomal insertions of the I-*Ppo*I protein variants were generated by micro-injecting embryos of the *An. gambiae* G3 laboratory strain. Of all the transgenic strains examined, ^gfp^124L-2, since renamed by Target Malaria as Ag(PMB)1 (for *An. gambiae* Paternal Male Bias strain 1), showed high sex ratio distortion among progeny of transgenic males (approximately 95% males), without significantly affecting male fertility and fitness [11]. Ag(PMB)1 contains a transgene that encodes a variant of I-*Ppo*I with an amino acid substitution at position 124 in its dimerization domain and which is fused to eGFP with the F2A self-cleaving peptide (eGFP(F2A)I-*Ppo*I-124L). This integrated into chromosome 3R-36D in position 47,762,151bp, from where the sex-distortion phenotype was stably inherited over consecutive generations. In large cage experiments, weekly inoculative releases of transgenic Ag(PMB)1 males led to a reduction in frequency of females and egg productivity of the population over successive generations consistent with model predictions [23].

**Figure 1.**
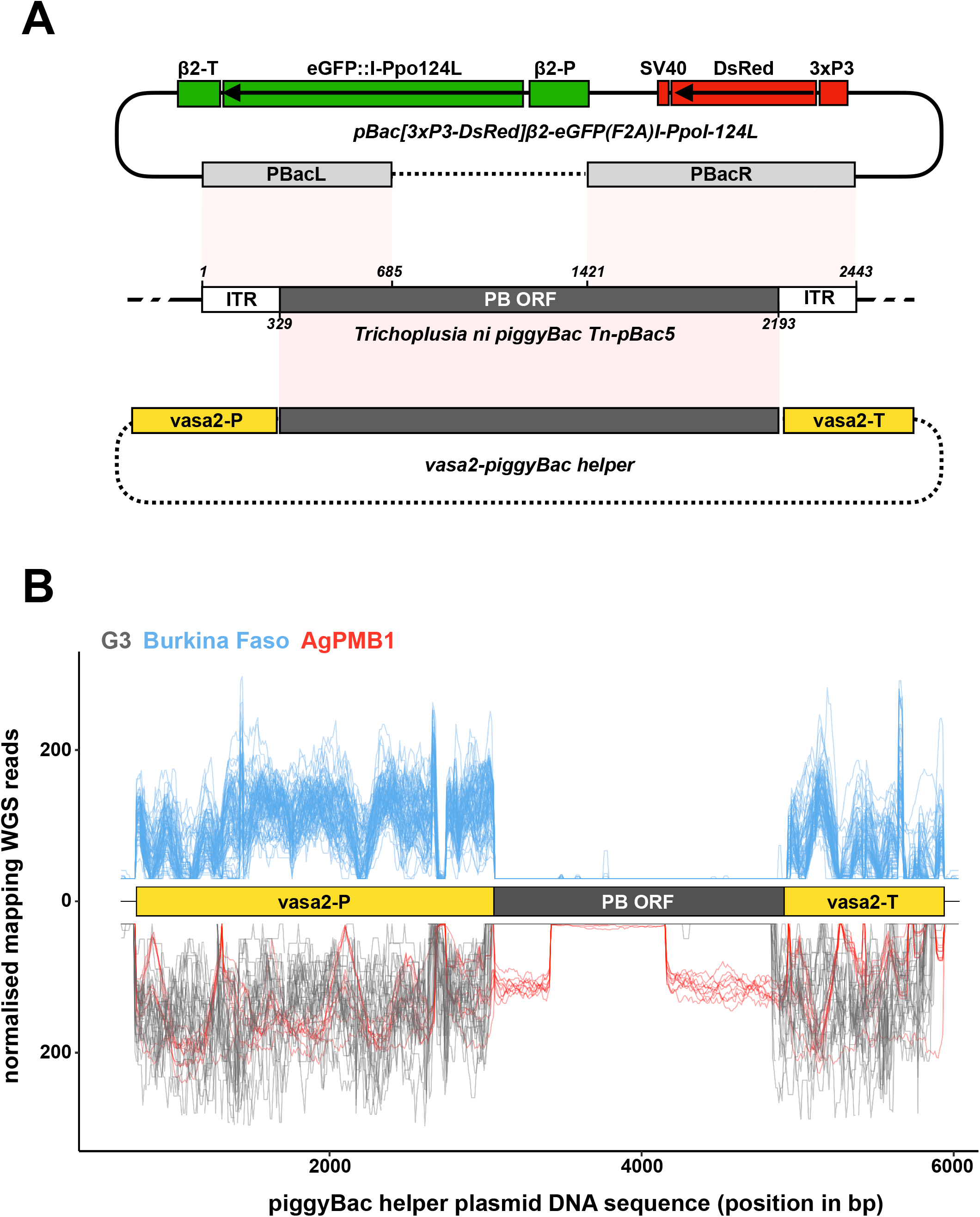
piggyBac transposase components in laboratory and wild type individuals. **(A)** Schematic of a wild type piggyBac (PB) transposon from *Trichoplusia ni* (middle; NCBI accession DQ236240.1), the Ag(PMB)1 transformation construct (top) and the PB helper plasmid (bottom). Shown are the regions of the endogenous PB locus present in the two microinjected plasmids highlighting how both the transformation and helper plasmid lack the complete machinery required for transposon mobility. pBacL (left) and pBacR (right) arms present in the pBac[3xP3-DsRed]β2-eGFP(F2A)I-*Ppo*I-124L transformation construct contain the entire flanking inverted terminal repeats (ITRs) and partial regions of the PB open reading frame (ORF). The helper PB plasmid containing the complete PB ORF driven from the *vasa*2 regulatory regions lacks the flanking ITRs. Sequences of the transformation construct are integrated in the genome are shown in solid lines, whereas sequences present only transiently in injected individuals are shown in dashed lines. **(B)** Mapping of whole genome sequencing reads from the G3 and Ag(PMB)1 controls (bottom in negative y-axis; grey and red) and the 81 wild type individuals collected in Burkina Faso villages (top; blue). The position of the *An. gambiae vasa* regulatory regions (yellow boxes) and the PB transposase ORF (black box) is also shown. Reads are normalized by scaling counts to the number of reads in the most abundant sample.

Since Ag(PMB)1 males are fertile their release could produce viable offspring in the field, unlike sterile males. This would provide invaluable information about how transgenic, laboratory-reared males of *An. gambiae* disperse spatially once released through detection of their offspring, even from small-scale releases aimed at capacity-building and methodology development. However, unlike fs-RIDL strains that were directly developed for deployment, the lack of a conditional expression system to control activity of eGFP(F2A)I-*Ppo*I-124L makes this strain unsuitable for large-scale programs aiming directly for population suppression. The Ag(PMB)1 transgene was not designed for gene drive in its current form - for example it is not able to home into targeted sequences. It also does not display any fitness advantage over wild type mosquitoes. Consistent with this, models have shown that the Ag(PMB)1 transgene would disappear over time when releases are discontinued [16].

However, one feature that is unique to an autosomal X-shredder is the possibility that it could move to the Y-chromosome after release. If active, a Y-chromosome linked X-shredder could directly benefit from the increased transmission of the Y-chromosome, thus increasing in frequency, persisting longer and dispersing further than initial planned. The sequential events required for such a driving Y to occur can be mapped to pathways to harm using a problem formulation approach adopted widely in environmental risk assessments (Supplementary Figure 1, Supplementary Figure 2). Three requirements must be fulfilled for a driving Y to occur: (1) the autosomal X-shredder must first move from its original autosomal position and become physically linked to the Y-chromosome; (2) the X-shredder would need to be expressed from its new position on the Y-chromosome during late spermatogenesis in a spatiotemporal manner that is similar to its original expression from the autosome; and (3) it should impart no significant cost to male fertility or male viability as a result of its new Y-chromosome-linkage (Supplementary Figure 1, Supplementary Figure 2).

With regard to requirement (1), there are two possible mechanisms that could result in an autosomal transgene moving to the Y-chromosome: (i) a transposase-mediated transposition to the Y-chromosome of the PB transposable element that was used to create the transgenic strain, or (ii) a recombination-mediated reciprocal translocation resulting in large chromosomal rearrangements between the autosome and the Y-chromosome. Of the two mechanisms, translocation is the less likely route, because translocations between autosomal segments and the Y-chromosome occur very rarely in nature (see Discussion). Furthermore, because these events are so rare, they are unlikely to be detected in small laboratory reared populations. On the other hand, transposition from the autosome to the Y through the remobilization of the PB transposon could be possible, if the X-shredder transgene co-occurs in a genome containing an active PB transposase.

In this paper, we addressed the possibility of transgene remobilization by examining whole genome sequence data generated from individuals of the Ag(PMB)1 strain, the wild type G3 laboratory colony, and field-derived wild-caught individuals from Burkina Faso for the presence of the PB transposase. With regard to requirement (2) above – expression from the Y – we generated two independent transgenic strains containing the eGFP(F2A)I-*Ppo*I-124L (the same in Ag(PMB)1) construct on the Y-chromosome and evaluated the level of expression and sex-ratio distortion. Finally, we discuss implications for male fertility depending on the route of movement to the Y-chromosome.

## Results

### Evaluating the remobilization potential of the autosomal Ag(PMB)1 X-shredder transgene

The Ag(PMB)1 strain was generated by Galizi et al. [11] by micro-injecting *An. gambiae* G3 embryos with a mixture of the transformation plasmid (pBac[3xP3-DsRed]β2-eGFP(F2A)I-*Ppo*I-124L) and a helper plasmid, containing the PB transposase expressed from the *vasa* regulatory regions [24], which direct expression in germline tissues (Figure 1A) [25]. By providing the PB transposase only in *trans* from a transiently co-injected helper plasmid, the transformation construct itself becomes immobilized once it is integrated in the genome. This is because, unlike the complete PB transposable element, the transgene lacks the complete transposase enzyme that is required for remobilization. Therefore, integrated PB transgenic constructs can only be remobilized in mosquito transgenic strains, if a PB transposase source is available [26].

To assess the stability of the Ag(PMB)1 transgene, we first evaluated whether the original insertion site, as described in Galizi et al. [11], has remained stable in chromosome 3R-36D at position 47,762,151 bp, over the approximately 100 generations since its initial generation. We designed PCR primers that span the PB transgene and genomic boundary as originally reported and repeated the PCR using genomic DNA from four pools of five transgenic larvae each of the Ag(PMB)1 strain. PCR amplicons were Sanger sequenced and blasted against the *An. gambiae* genome assembly. Compared to the inverse PCR results from Galizi et al. [11], we found that the genomic integration site has remained identical (Supplementary Figure 3). This suggests that the Ag(PMB)1 transgene has not remobilized in the strain, indicating that PB transposase does not occur naturally in the genome of the laboratory colony. It also suggests that none of the other naturally occurring transposable elements present in this strain are able to remobilize the Ag(PMB)1 transgene, in the absence of PB.

To confirm this result, we next tested whether we could detect the gene encoding PB transposase in the genomes of the G3 or Ag(PMB)1 strains. This would exclude the possibility that PB transposase gene is present but is either non-functional, e.g. through mutations in its open reading frame, or suppressed by gene silencing by piRNAs [27,28]. To do this, we generated whole genome sequence (WGS) libraries from genomic DNA extracted from 10 individuals (5 females, 5 males) of the Ag(PMB)1 strain and downloaded WGS libraries from 24 previously sequenced G3 individuals from the same insectary colony (PRJNA397539). We mapped the WGS data to the PB helper plasmid originally used to generate the Ag(PMB)1 transgenic strain, containing the PB transposase driven by the 5’ and 3’ regulatory regions of *An. gambiae vasa* gene. Mapping WGS reads against the helper plasmid ensured that the coding sequence evaluated is experimentally verified to catalyze excision of PB transgenes, instead of a different transposable element that may be related at the sequence level but is unable to excise PB transgenes. Conveniently, the helper plasmid included internal positive controls, in the form of regulatory sequences from the endogenous single-copy *vasa* gene and parts of the flanking PB left and right arms of the Ag(PMB)1 transgene (Figure 1A). We observed a high number of mapping WGS reads from G3 samples against the *vasa*-derived regulatory sequences on the helper plasmid, but no continuous mapping in the region corresponding to the PB transposase enzyme (Figure 1B). For the Ag(PMB)1 stain, genomic reads mapped to both the endogenous *vasa* regulatory sequences and to internal sequences of the PB ORF that correspond with the parts of PB left and right arms included in the transformation construct used to generate Ag(PMB)1, as expected (Figure 1A). No reads were detected on the PB coding sequence that is excluded in the transformation construct, thus making it non-autonomous (Figure 1A and B). We then repeated the same analysis using single-mosquito WGS data from 81 field-caught individuals collected in Burkina Faso in 2012 (PRJEB1670), which is the site of previous releases of genetically sterile male mosquitoes (Supplementary Figure 4)[29]. Similar to the results from the G3 samples, no reads mapped to the part of the helper plasmid encoding the PB transposase open reading frame with reads mapping exclusively to the regions of the endogenous *vasa* gene. Together, these results suggest that the PB transposase is unlikely to be in the same background as the autosomal Ag(PMB)1 transgene that is being considered for release.

### X-shredder expression from the Y-chromosome during spermatogenesis

The second requirement for the Ag(PMB)1 X-shredder to display gene drive and invasiveness, assuming the transgene has first moved to the Y-chromosome, is that it is expressed in a correct spatiotemporal manner from its new location. In the Ag(PMB)1 strain, X-shredding is achieved through the expression of the eGFP(F2A)I-*Ppo*I-124L transgene from the *An. gambiae beta2-tubulin* regulatory regions, which is active shortly before the first meiotic division in primary spermatocytes and continues throughout the subsequent stages of spermatozoa differentiation [30]. In previous work, we have shown that transgenes driven from this promoter are strongly expressed when located on *An. gambiae* autosomes, but when they are inserted on the X-chromosome expression is undetectable [31]. This includes various X-chromosome-linked X-shredder variants, where no significant expression or sex bias was observed [11]. Similar observations of X-linked transgene transcriptional suppression around meiosis have been made in other species as well [32–34]. This phenomenon, called meiotic sex chromosome inactivation (MSCI), is thought to be one of the main driving forces leading to the observed paucity of sperm-specific genes on the X chromosome, both in *An. gambiae* mosquitoes and in other species [35–37]. By comparison, much less is known about transgene expression during spermatogenesis from the *An. gambiae* Y chromosome, which is estimated to be around 26 Mbp long, approximately 10% of the mosquito genome [38], and is composed nearly entirely of a few massively amplified, tandemly arrayed repeats and five known genes [39].

To confirm whether MSCI has a similarly inhibitory effect on transgene expression during spermatogenesis on the Y-chromosome as the X-chromosome, we generated two independent transgenic strains harboring the Ag(PMB)1 transgene, eGFP(F2A)I-*Ppo*I-124L, on the *An. gambiae* Y-chromosome. The first transgenic strain, called YpBac-β2-^gfp^124L, was generated by random PB integration. We sequenced the insertion site of the YpBac-β2-^gfp^124L transgene by inverse PCR on genomic data extracted from transgenic males and found that the construct had inserted within the ubiquitous Y-chromosome-specific transposable element *zanzibar* [40] (Figure 2A). The second transgenic strain, called YattP-β2-^gfp^124L, was obtained by secondary φC31 site-specific integration into an AttP docking site we previously inserted on the Y-chromosome [41]. Similar to the YpBac-β2-^gfp^124L strain, the AttP site is located in a region of the Y-chromosome containing the *zanzibar* repeat, though it is not possible to estimate the distance between these two insertions given the lack of a continuous Y-chromosome genome assembly [40].

**Figure 2.**
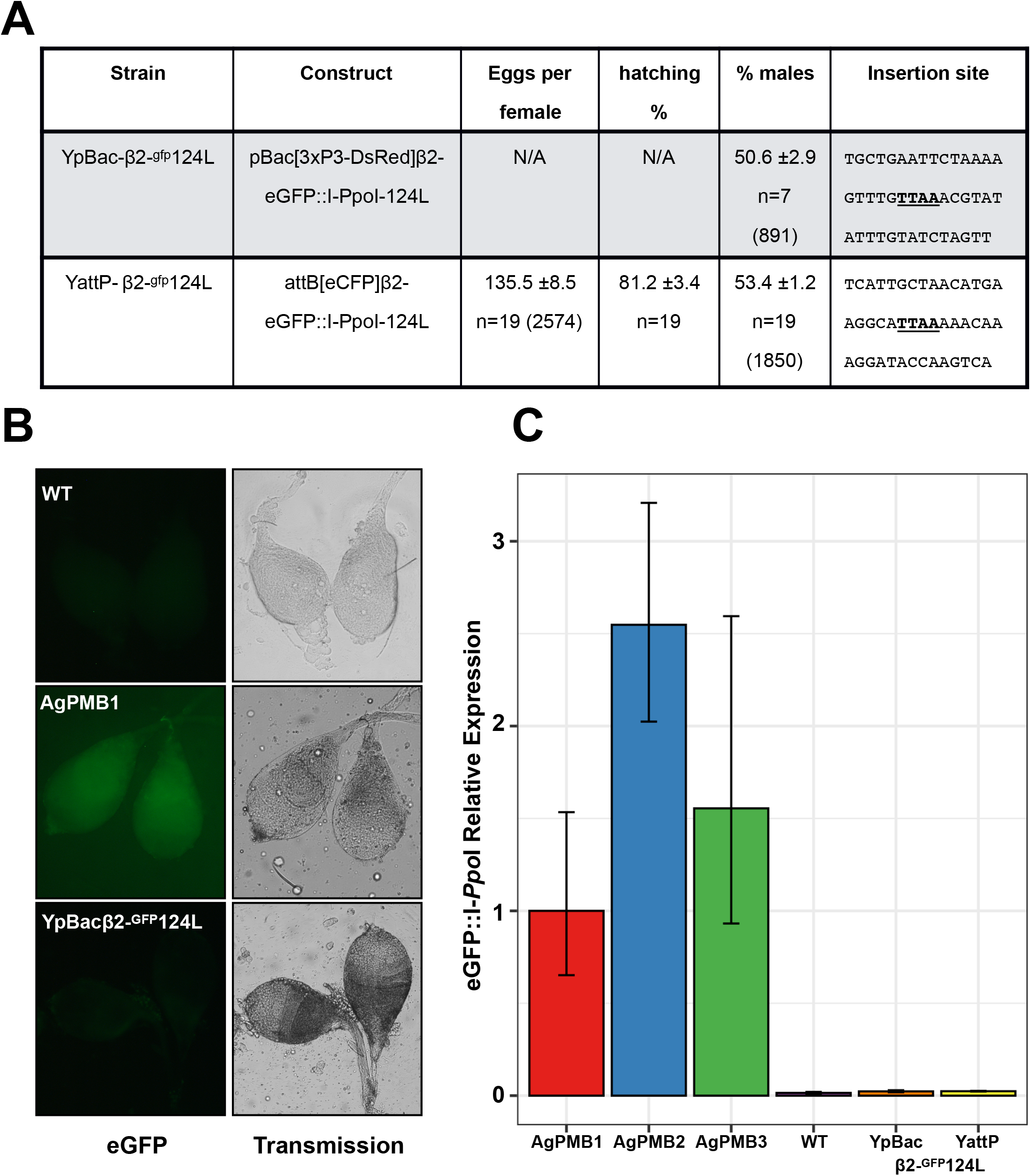
Transcriptional suppression of Y-linked X-shredder constructs abolishes sex ratio distortion. **(A)** eGFP fluorescence from dissected wild type (WT), Ag(PMB)1 and YpBac-β2-^gfp^124L testis. **(B)** Quantitative RT-PCR showing the relative expression of eGFP(F2A)I-*Ppo*I variants in different X-shredder strains, compared to expression in the Ag(PMB)1 strain. Expression of the X-shredder is undetectable in both Y-chromosome insertions. Expression levels from two additional autosomal strains, Ag(PMB)2 and Ag(PMB)3, which also led to sex ratio distortion are shown. **(C)** Progeny analysis of males from the two Y linked X-shredder strains crossed to wild-type females. Shown is the average number of eggs laid per n females analyzed (± represents the standard error of the mean; SEM). Average percentage of larvae hatching from the eggs (±SEM), from n females analyzed. Average percentage of males in the progeny (±SEM) from n females. The total number of eggs or individuals counted in each experiment is given in parentheses. Sequences (20bp each side) flanking the PB integration site (TTAA) of the transformation constructs are also shown.

As would be expected from Y-linked insertions, transgenic offspring from males of both Y-linked strains and wild-type G3 females were exclusively male. Testes from transgenic males of both strains displayed no detectable eGFP signal by fluorescence microscopy, which would be expected if the eGFP(F2A)I-*Ppo*I-124L X-shredder transgene was expressed (Figure 2B). Conversely, expression from the 3xP3-DsRed transformation markers in both strains was phenotypically indistinguishable from autosomal insertions (data not shown), suggesting that Y-chromosome linkage does not interfere with somatic expression of transgenes, at least from these two positions [41]. We next analyzed the levels of eGFP(F2A)I-*Ppo*I-124L transcription in the testes of the YattP-β2-^gfp^124L and YpBac-β2-^gfp^124L strains, compared to the Ag(PMB)1 strain. As a control, we also evaluated expression in testes of two additional strains from the Galizi et al. [11] study (^gfp^124L-3 and ^gfp^111A-2, called here Ag(PMB)2 and Ag(PMB)3, respectively) and wild-type male. Results from the quantitative RT–PCRs show no significant levels of eGFP(F2A)I-*Ppo*I-124-L expression in the testes from both Y-linked transgenic lines (Figure 2C). Consistent with the lack of X-shredder expression, we did not detect any sex bias among progeny when transgenic males from each Y-linked strain were crossed to wild type G3 females (Figure 2A). These results highlight that, as would be expected from MSCI, the Y-chromosome is not permissive to transgene expression from the *beta2-tubulin* promoter during late spermatogenesis, similarly to the X-chromosome.

## Discussion

Genetic control strategies that aim to suppressing wild populations of mosquito disease vectors have garnered significant interest, and field trials of a number of these systems, including classical SIT, IIT *(Wolbachia-based* sterility), and RIDL are now underway [42,43]. Synthetic sex-ratio distorters based on X-chromosome shredding have now been developed in *An. gambiae* [11,44] and more recently in *Dr. melanogaster* [45]. This system has been shown both theoretically and experimentally to be more efficient than classical SIT, in terms of the number of insects that need to be released [11,15,16]. In their most basic form, autosomal X-shredder constructs are designed to be self-limiting, and their release can potentially result in local and limited suppression if sufficient males are released over a long enough period. Nonetheless, field releases of fertile autosomal X-shredder males have not yet been conducted.

Here, we have evaluated the theoretical possibility whereby a released autosomal X-shredder could convert into a self-sustaining, driving Y-chromosome after release. The requirements for such an event to occur include: (1) movement of the X-shredder to the Y-chromosome; (2) its subsequent expression from the Y during late stages of spermatogenesis; and, (3) no cost to male fitness. We reason that there are two possible mechanisms that could result in the linkage of the autosomal Ag(PMB)1 to the Y-chromosome: (1) a transposase-mediated remobilization of the transgene and (2) a large chromosomal rearrangement resulting in the reciprocal translocation between the region of chromosome 3R containing the transgene and the Y-chromosome.

We have evaluated the potential of remobilization of the Ag(PMB)1 transgenic construct through transposition, mediated by the intact PB inverted terminal repeats (ITRs) on either side of the transgene cassettes, which were used for the initial generation of the strain. Their presence makes it at least theoretically possible that the Ag(PMB)1 transgene could remobilize from its position on chromosome 3R, if a source of the PB transposase occurs in *trans.* We therefore evaluated whether the PB transposase is present in the genomes of the laboratory Ag(PMB)1 and G3 colonies, and also in field-derived samples from Burkina Faso. We found no evidence of the complete PB transposase coding sequence in any of the samples we sequenced, suggesting that PB is not present at appreciable frequencies in *An. gambiae* mosquitoes sampled from nature. This result is supported by the long-term stability of the Ag(PMB)1 insertion site over 100 generations since its original construction, a result that suggests that other naturally occurring repetitive elements in the genome of the Ag(PMB)1 are not functionally capable of PB transgene remobilization.

We were not able to test the second possibility of remobilization by chromosomal translocation, as these occur very rarely during meiosis. Under standard laboratory rearing of the Ag(PMB)1 strain in over 5 years, translocations involving the autosomal transgene and the Y-chromosome have never been detected – an event that would be immediately noticeable since fluorescent transgenic individuals would only be male. This extends to scaled experimental conditions, in which large numbers of Ag(PMB)1 individuals were screened by high-throughput sorting of individuals using the COPAS sorter based on the 3X-P3-DsRed marker and then subsequently separated by sex at the pupal stage [23]. This is, in part, expected because of how rarely such events occur, meaning that experimentally verified rates for such events are not readily available for the Ag(PMB)1 strain. When translocations between autosomes and the Y are desired they can be artificially induced in the laboratory, for example with ionizing radiation, chemical agents, or UV radiation. This is commonly done for insects to link selectable phenotypes to the Y-chromosome, in so-called genetic sexing strains (GSSs). GSSs are developed so that males can be separated from females on a large scale in insect bio-factories that produce animals for genetic control programs, such as SIT [46]. This is done by linking selectable traits, for example insect color or high temperature tolerance, to the Y-chromosome using induced reciprocal translocations of mutant alleles located on autosomes. Once generated, these GSSs are then maintained in large numbers, with billions of insects being produced weekly. The large colony size makes it possible to detect rare events that result in the breakdown of linkage between maleness and selectable trait. One of two ways this can happen is through a “reverse” reciprocal translocation involving the previously modified Y-chromosome and an autosome (known as a type 2 recombination event) [47]. Because such events lead to a breakdown of the genetic sexing system and restore male fertility of the semi-sterile males (arising from the translocation itself) leading to their accumulation, their occurrence is tightly monitored in large-scale rearing operations. In the only report that quantified the rate of type-2 recombination and distinguished it from type-1 (which does not involve the translocated Y-chromosome and is more common among the two) in a GSS of the Mediterranean fruitfly *Ceratitis capitata*, the rate was estimated to be 10^-5^ or less, i.e. occurring in less than 1 out of 100,000 male individuals [47]. In this case however, there are two significant factors that would indicate that the rate of an initial, uninduced autosome-Y translocation would be much lower. First, the rate of recombination in the GSS describes an event reversing a previously induced autosome-Y translocation, which is likely to be largely mediated by homologous sequences that are now present on the two translocated Y fragments; a homology that does not normally exist between autosomes and the Y chromosome. Therefore, the expected recombination rate resulting in a reciprocal translocation between an autosome and Y would be lower. Second, the reversion of the previously translocated autosomal fragment on the Y restores male fertility, that was first compromised by the translocation to the Y (because of gamete chromosomal imbalance – discussed below) [47]. This means that type-2 recombinant males will have more viable offspring increasing their rate of occurrence in the population. Together, these factors suggests that the probability of a translocation event involving the autosomal Ag(PMB)1 transgene and the Y-chromosome in progeny of released males born in the field is much lower than 10^-5^. This rate would depend on the size of the Y-chromosome and the relative rate of recombination in the male germline, which in *An. gambiae* is approximately 1.6 cM Mb^-1^ for chromosome 3R [48] and similar between males and females [49]. Finally, it is worth noting that since there is no recombination in *Drosophila melanogaster* males, data on autosome-Y translocation frequencies do not exist in this insect model, to our knowledge.

In the unlikely event that the transgene was to move to the Y-chromosome, we provide data regarding the expression of the X-shredder from this chromosome, and conclude that MSCI during spermatogenesis does affect the Y-chromosome of *An. gambiae.* Our results from two transgenic strains harboring the Ag(PMB)1 X-shredder transgene in two different positions on the Y-chromosome, reveal transcriptional suppression during late spermatogenesis from the *beta2-tubulin* promoter, complementing our previous work which confirmed this for the *An. gambiae* X-chromosome [11,35,36]. We found no evidence of X-shredder expression by RT-PCR, nor by fluorescence microscopy of transgenic testis. Offspring of transgenic males from both Y-linked strains therefore had sex-ratios similar to wild-type males. Hence, even if the Ag(PMB)1 transgene successfully moved to the Y-chromosome by transposition it is unlikely that the X-shedder would be active. Since MSCI-factors regulating transcriptional suppression physically spread across the sex chromosomes after becoming localized on their unsynapsed axes [50,51], it is also expected that translocated autosomal fragments would become suppressed by MSCI during meiotic stages of spermatogenesis. Therefore, the weight of evidence argues strongly against the likelihood of movement of the Ag(PMB)1 transgene to the Y chromosome, particularly via transposition. However, the equally necessary perquisite for a pathway to a driving Y, namely expression of the X-shredder on the Y chromosome during male meiosis, seems highly implausible based on the evidence presented here.

The final requirement for a Y-linked X-shredder to spread through populations is that its movement to the Y-chromosome and subsequent expression from it would have no significant fitness costs to males harboring it. Such fitness costs would counteract the theoretical advantage gained by the Y-linked X-shredder from increased transmission through elimination of X-bearing sperm. Among the factors determining these fitness costs, the largest contributors would likely be the mechanism leading to Y-linkage and the outcomes of this movement on each chromosome. Reciprocal translocations between an autosome, in this case 3R, and the Y-chromosome typically result in significant male fertility costs [52]. Because of the simultaneous segregation of non-homologous centromeres (adjacent-1 segregation) during meiosis, only 50% of the offspring produced by males are genetically balanced, i.e. males are 50% sterile [53]. In certain cases, this semi-sterility can be even higher, for example in an *An. arabiensis* GSS showing 73.3% male sterility [53]. Therefore, a Y-linked X-shredder that arose through a translocation event would likely display such high male fertility costs that it would rapidly disappear from the population. For transposition-mediated Y-linkage male fitness costs cannot be predicted *a priori*, as gamete balance and genic content would depend on both the excision event (i.e. how much of the surrounding chromosome is excised) and on the integration position on the Y-chromosome (i.e. subsequent knock-out of genes essential for male fitness such as the male-determining gene).

In summary, the findings of the current study support the low probability of Y-chromosome linkage through remobilization of the transposon used for the initial genetic transformation. Moreover, even if such as rare event occurred, where the X-shredder would become linked to Y-chromosome, activity of the X-shredder at the required stage of spermatogenesis would likely be impeded via chromosome wide suppression of gene expression on sex chromosomes. Prospects for the systematic generation of an X-shredding-based Y-drive for mosquito control in the future, will need to find ways to circumvent this transcriptional suppression, for example using alternative germline specific promoters [37].

## Methods

### Mosquito rearing

Wild-type *An. gambiae* strain (G3) and transgenic mosquito strains were reared under standard conditions at 28 °C and 80% relative humidity with access to fish food as larvae and 5% (wt/vol) glucose solution as adults. For egg production, young adult mosquitoes (3–5 d after emergence) were allowed to mate for at least 6 d and then fed on mice. Three days later, an egg bowl containing rearing water (dH2O supplemented with 0.1% pure salt) was placed in the cage. One to two days after hatching, the larvae (L1 stage) were placed into rearing water containing trays.

### Assaying transgene chromosomal location

Primers were designed to amplify across the 5’ and 3’ genomic insertion boundaries of the Ag(PMB)1 transgene. DNA was extracted from 4 individual pools (containing 5 transgenic larvae each) of Ag(PMB)1 mosquitoes following >100 generations of lab maintenance. PCR across the 5’ and 3’ transgene flanking regions was performed using: pB-5SEQ (5’-CGCGCTATTTAGAAAGAGAGA) with genomic primer RH57 (5’-GAAAACCTTACAAGCGTCTTCAA) to amplify the 5’ junction and pB-3SEQ (5’-CGATAAAACACATGCGTCAATT) with RH53 (5’-GATATTATCGCGCTCGGTCC) to amplify the 3’ junction. Each PCR amplicon was Sanger sequenced using both forward and reverse primers and the resulting sequences were aligned to compare genomic flanking and transgenic sequences to the Galizi 2014 data [11].

### Mosquito whole genome sequencing and read mapping

*Anopheles gambiae* WGS reads from 81 individuals collected in Burkina Faso in 2012 were downloaded from the European Nucleotide Archive (Study Accession: PRJEB1670; Supplementary Table 1). WGS data from the G3 laboratory colony were downloaded from the SRA (Bioproject Accession: PRJNA397539). Genomic DNA from 10 Ag(PMB)1 individuals was extracted using the Blood and Tissue Kit (Qiagen). For each sample, 100ng of input gDNA was sheared using Covaris for a 350bp insert size. Library preparation was performed using the Illumina TruSeq Nano kit. Each sample was tagged with a unique barcode, followed by three 2×150bp High Output V2.5 paired-end sequencing runs on the Illumina NextSeq550 platform, obtaining an average of 265M reads per sample. WGS data from the Ag(PBM)1 have been deposited and will be made available for submission of this manuscript. Fastq reads were quality checked with FastQC [54] and converted to fasta format. Reads were then mapped against the *vasa* driven piggyBac plasmid [24] using blast blast-2.2.26/bin/blastall -i db.fa -d sample.fasta -p blastn -F “m L” -U T -e 1-e4 -a 40 -v 5 -b 40000 -K 40000. Only alignments with 98% identity over the entire read length were kept. Coverage was computed for each sample and normalized to the read depth of the most deeply sequenced sample using the following formula Xi = Xi/(Xi/Xmax). To clarify plotting, read depth is reported every 10 bp.

### Generation of Y-chromosome linked X-shredder transgenic strains

The YpBac-β2-^gfp^124L transgenic strain was generated as described in Galizi et al. [11]. Briefly, *An. gambiae* G3 embryos were injected with a mixture of 0.2 μg μl-1 of the pBac[3xP3-DsRed]β2-eGFP(F2A)l-*Ppol*-124L plasmid and 0.4 μg μl-1 of helper plasmid containing a *vasa*-driven piggyBac transposase [24]. The hatched larvae were screened for transient expression of the DsRed marker and positives (~54%) crossed to wild-type mosquitoes. F1 progeny were analysed for DsRed fluorescence and positives were crossed individually with wild-type mosquitoes to obtain transgenic lines. The transgene of one strain derived from a G0 male was identified that was transmitted exclusively to F1 sons, indicating Y-chromosome integration. The stain, now called YpBac-β2-^gfp^124L was established and maintained by crossing to wild type females. The YattP-β2-^gfp^124L strain was generated by co-injecting the pBac[3xP3-DsRed]β2-eGFP(F2A)I-*Ppo*I-124L construct with a vasa2-driven ΦC31 integrase helper plasmid [24] into eggs of a strain containing a Y-chromosome AttP docking site [41]. Crosses and screening were performed as above.

### Sex ratio and fertility assays

To assay adult sex ratio, 10 transgenic males of each line were crossed to 10 wild-type females. In all crosses, mosquitoes were allowed to mate for 3-5 days after the blood meal gravid females were placed individually in oviposition cups. Larvae were reared to adulthood and sex was counted. The number of eggs laid as well as the number of larvae hatching were also counted, but only for the YattP-β2-^gfp^124L to assay male fertility.

### qRT-PCR analysis

qRT-PCRs were performed on mosquito total RNA as described in Galizi et al. [11]. Briefly, 10 pairs of testes from each transgenic strain were pooled to constitute a biological replicate for total RNA and protein extraction using TRI reagent (Ambion). RNA was reverse-transcribed using Superscript II (Invitrogen) after TURBO DNA-free (Ambion) treatment following the manufacturer’s instructions. Quantitative real-time–PCRs (qRT–PCR) analyses were performed on cDNA using the Fast SYBR-Green master mix on a StepOnePlus system (Applied Biosystems).

Ribosomal protein Rpl19 gene was used for normalization. At least two independent biological replicates from independent crosses were subjected to duplicate technical assays. We used primers RPL19Fwd (5’-CCAACTCGCGACAAAACATTC-3’), RPL19Rev (5’-ACCGGCTTCTTGATGATCAGA-3’), eGFP-F (5’-CGGCGTGCAGTGCTTCA-3’) and eGFP-R (5’-CGGCGCGGGTCTTGT-3’).

## Acknowledgements

The authors would like to thank Kostas Bourtzis, Austin Burt, Andrea Crisanti and Samantha O’Loughlin for helpful discussions and support. This study was supported by the Italian Ministry of Education, University and Research (MIUR—D.M. no. 79 04.02.2014) grant to P.A.P. This study was also supported by a grant from the Foundation for the National Institutes of Health through the Vector-Based Control of Transmission: Discovery Research (VCTR) program of the Grand Challenges in Global Health initiative of the Bill & Melinda Gates Foundation. This research was supported by Research Grant No. IS-5180-19 from BARD, the United States – Israel Binational Agricultural Research and Development Fund to P.A.P. This research was supported by Research Grant No. 2388/19 from ISF, the Israel Science Foundation to P.A.P.

---

**Supplementary Figure 1.**
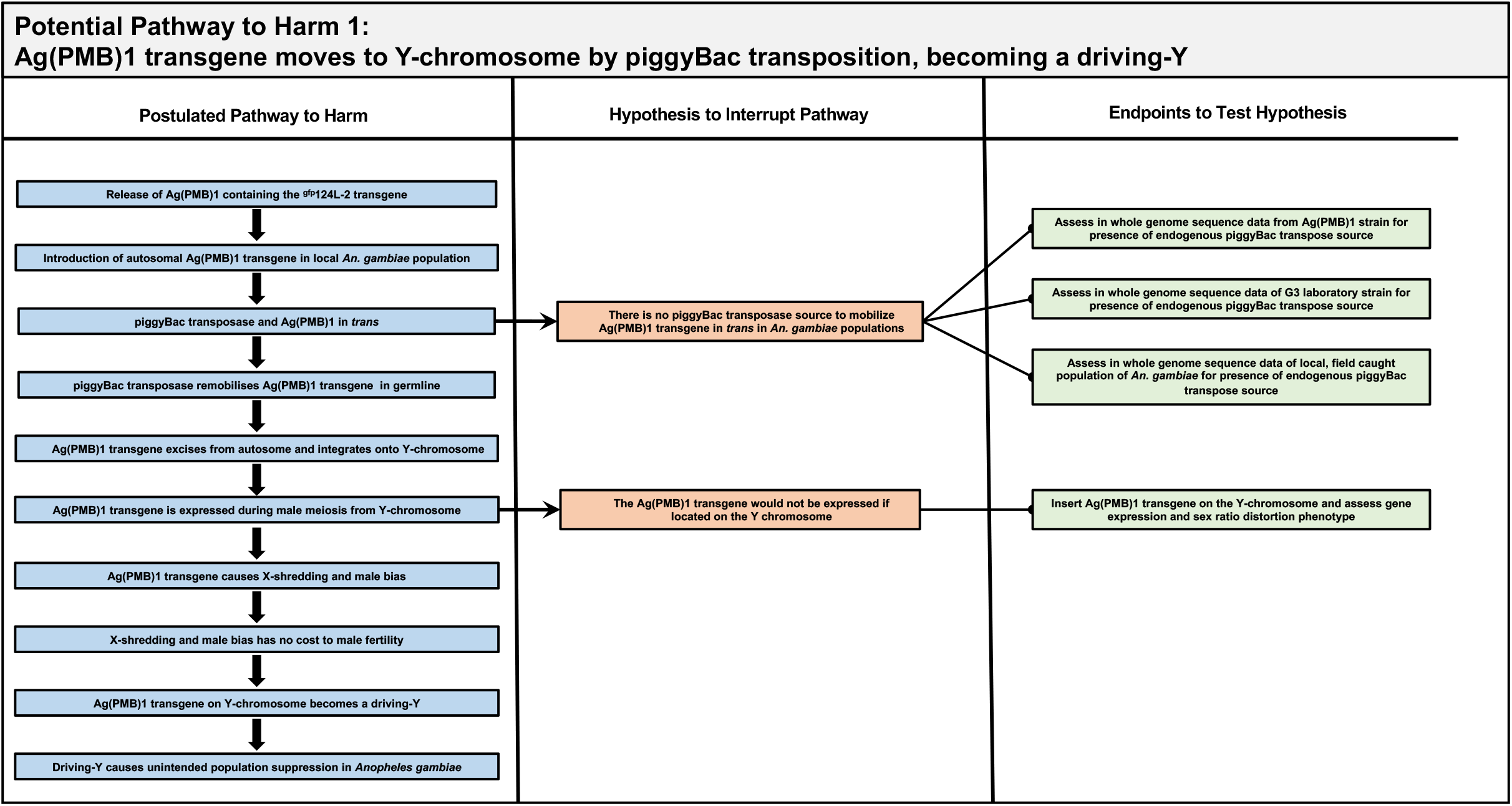
Potential Pathways to harm by piggyBac transposition of Ag(PMB)1 to the Y chromosome.

**Supplementary Figure 2.**
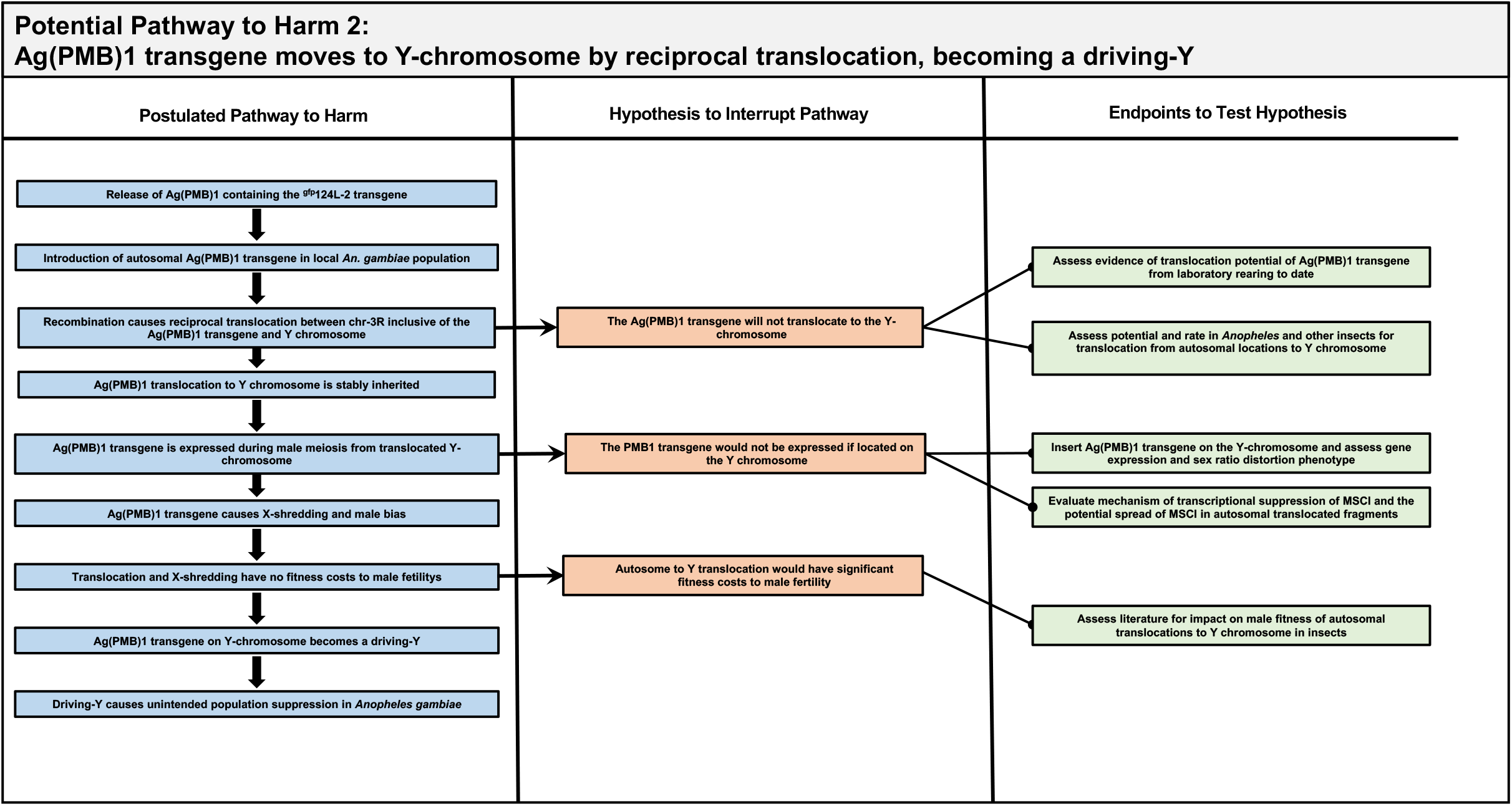
Potential Pathways to harm by reciprocal translocation of the autosomal Ag(PMB)1 to the Y chromosome.

**Supplementary Figure 3.**
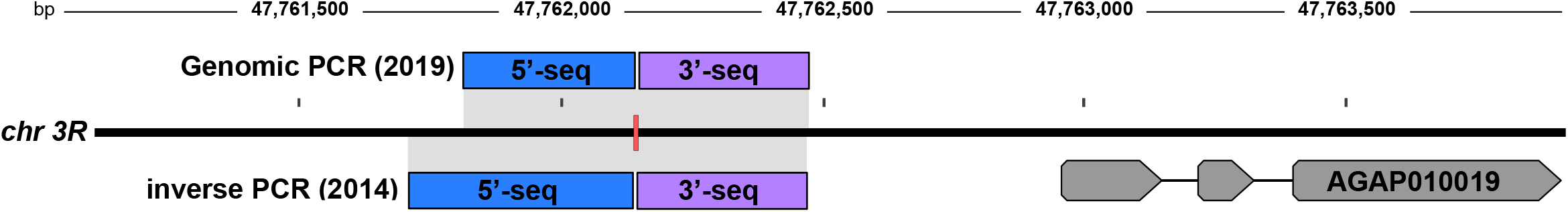
Analysis of Ag(PMB)1 transgene insertion. Blue and violet filled blocks highlight the 5’ and 3’ genomic sequences flanking the transgene based on sanger sequencing of the genomic PCRs. Shown are the results from the Galizi et al.[11] inverse-PCR analysis (below) and the sequencing of genomic PCRs from the present study (above). The location of Ag(PBM1) transgene is shown in a red box and is located on chromosomal arm 3R upstream of AGAP010019 gene (grey filled arrow). Base-pair positions on chromosome 3R are also shown. The analysis confirms that the insertion site of Ag(PMB)1 strain has remained stable over 100 generations of maintenance in the laboratory.

**Supplementary Figure 4.**
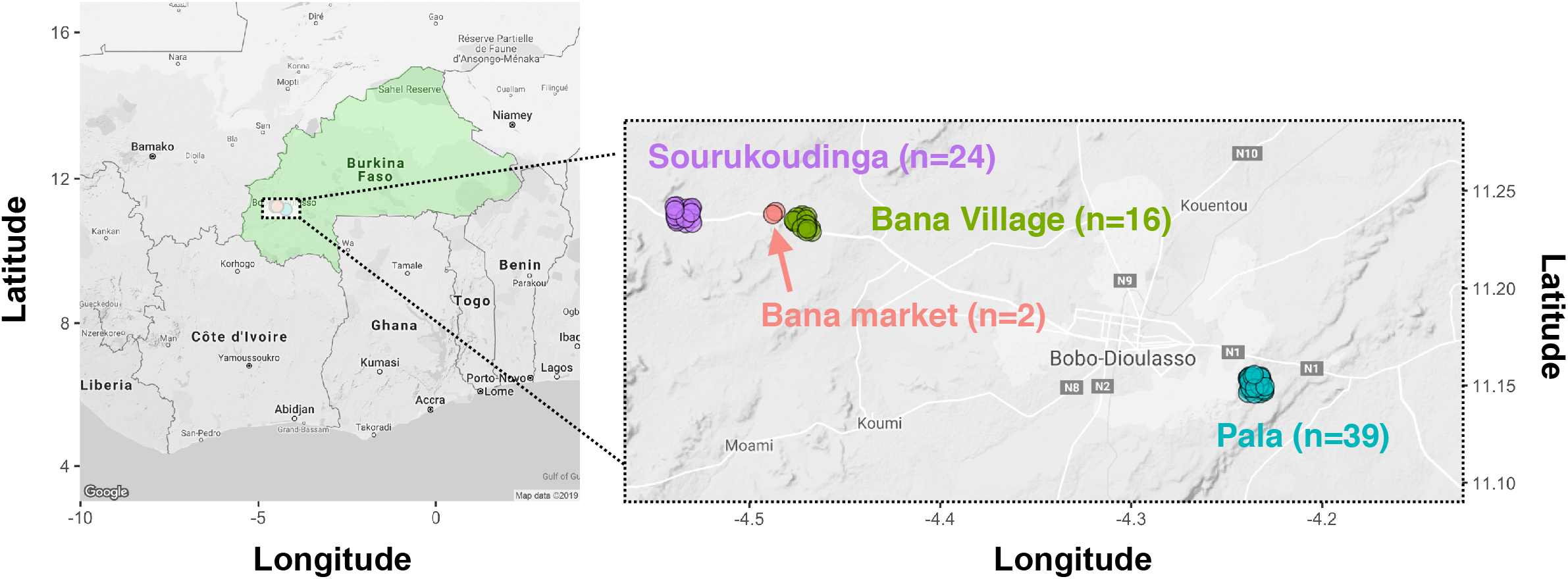
Location of Burkina Faso individuals used in this study. Longitude and latitude of samples collected in 4 locations in Burkina Faso in 2012. Names of villages are shown as well as the number of individuals from each site.

**Supplementary Table 1. Burkina Faso individuals Ag1000gs supplementary information**

